# Sex Differences in B2 SINE RNA Expression and its Role in Hippocampal Development

**DOI:** 10.1101/2025.11.18.689111

**Authors:** Troy A. Richter, Andrew A. Bartlett, Hannah E. Lapp, Erin T O’Neil, Ellie K Pritchard, Guia Guffanti, Susan L Zup, Richard G Hunter

**Affiliations:** Department of Psychology, University of Massachusetts, Boston, MA; Department of Psychiatry, McLean Hospital – Harvard Medical School, Belmont, MA; Department of Psychology, University of Texas at Austin, TX

**Keywords:** SINE, Sex, Brain, Hippocampus, Housekeeping gene

## Abstract

Once dismissed as “junk”, transposable elements (TE) have recently gained recognition for their regulatory roles, notably in the brain and in development. The brain is hormone-responsive and the hippocampus in particular is sensitive to circulating gonadal hormones. While transcriptionally active, TE function remains poorly understood, especially in the brain. We and others have shown that one particular TE RNA, B2 SINE ncRNA, is a regulator in the rodent hippocampus, especially after a psychologically stressful event like acute restraint stress. It is unknown, however, if B2 SINE ncRNA is necessary for the proper development of hippocampal neurons, and furthermore, if there is a sex difference in this development. This work investigates the difference in expression of B2 SINE RNA across sexes and its role in the development of primary hippocampal neurons. We utilized pooled locked nucleic acid (LNA) GapmeRs to knock down the expression of B2 SINE RNA and treated primary hippocampal neurons with dihydrotestosterone (DHT) to test if there is a difference in dendritic complexity. We used Sholl analysis to quantify the branching, number of tips, and Sholl mean. We found a sex difference in both B2 SINE, higher in males compared to females, and ß-actin, lower in males compared to females. Additionally, knocking down B2 SINE RNA results in a reduction of dendritic complexity in male but not in female neurons. Taken together, this work suggests that B2 SINE RNA is expressed differentially and plays an important role in the proper development of hippocampal neurons in a sex dependent manner. Our findings support the identification of a sex-specific biomarker that may enable individualized treatment of conditions influenced by sex. This is the first evidence of the role a transposable element’s RNA has in the regulation of the development of neurons and the first to show that differential regulation by sex.

**Summary Paragraph:** Long considered “junk DNA”, transposable elements have gained recognition as regulators in the genome. Some of these elements are transcriptionally active and have been known to control the expression of downstream protein coding genes. Recently, murine B2 SINE RNA has been shown to regulate glucocorticoid responsive genes in the rodent hippocampus. However, it is unknown if B2 SINE RNA is necessary for the proper development of hippocampal neuron structure and if there is a sex difference in this development. Here we show that B2 SINE is higher in the male hippocampus compared to females, and knocking down B2 SINE RNA results in a reduction of dendritic complexity in male but not in female neurons. This work shows that B2 SINE RNA is expressed differentially across sexes, and that this noncoding RNA plays an important role in the development of hippocampal neurons. These findings are the first to demonstrate the role for a transposable element’s RNA in the development of neuronal structure and the first to show that this is differential across sex. These findings support the identification of a sex specific biomarker that could lead to individual treatments of neurological disorders influenced by sex.

## Introduction

In most mammals, androgens are synthesized in the brain, gonads, and adrenal glands in both sexes and impart physiologically important consequences on the structure and function of the central nervous system. These consequences may lead to the development of neurodevelopmental and other neurological disorders like autism spectrum disorder, ADHD, Alzheimer’s disease, and schizophrenia, which all have different incidences across sexes ^1,2^. In terms of hippocampal synaptic plasticity, there are a number of sex differences already discovered. For example, after a stressful stimulus, males show a higher dendritic spine density in the CA1 region of the hippocampus compared to females ^3^. Much of the literature has focused on spine numbers, however, with fewer studies examining the development of overall dendritic complexity^4–6^. Receptors for androgens (AR) are present at high levels in the nuclei of hippocampal neurons as well as extranuclear sites like the plasma membrane, mitochondria, and synaptic vesicles ^7–11^. The presence of AR in the hippocampus suggests that the actions of androgens, play a role in the function of the hippocampus itself in adulthood ^12^. However, little attention has been paid to the developmental role of androgens in organizing fine structure of hippocampal neurons.

The classic method of action for activated AR is to form a dimer and translocate to the nucleus, where it acts as a transcription factor, promoting or suppressing the transcription of androgen-responsive genes. In this way, circulating androgens can interact with the genome and confer changes to the cell. Studies have shown that steroids are synthesized in the developing hippocampus and throughout life ^13^. Furthermore, these synthesized hormones in the hippocampus can contribute to the modulation of memory processes, especially if the levels of these hormones are altered during development ^14–17^. While studies are increasingly focused on how sex hormones can alter synaptic plasticity, specifically at the level of synapses and dendritic spines, there are currently no studies exploring the sex differences that may change the dendritic arborization of primary hippocampal neurons after transient treatment with dihydrotestosterone (DHT).

Transposable elements (TEs) make up the largest fraction of the mammalian genome estimated to occupy anywhere from just over half to two-thirds of a given genome ^18–20^. Historically viewed as “junk” DNA, the ENCODE project found that nearly 80% of the human genome was transcriptionally active in at least one cell type and that much of this was TE-derived RNA ^21^. TE expression is tightly regulated, tissue-specific and in some contexts necessary for host biological function ^22^. Given their genomic occupancy and activity in eukaryotes, TEs represent a largely unexplored class of regulatory elements. Murine SINEs are short (<300bp) repetitive elements with the subclass B2 being the most common in rodents. These elements are transcribed into noncoding RNA by RNA Polymerase III. The most well-known event by which B2 SINEs are transcribed into RNA is during cellular heat shock. In response to heat shock, B2 SINE RNA is a master regulator of the cellular response. The heat shock response involves the rapid coordination of cellular machinery simultaneously upregulating heat shock proteins while downregulating other processes including the expression of RNA Pol II-dependent HKG expression for example ß-actin ^23^. RNA Pol III-dependent expression of HKGs, for example 7SK, remain unchanged during heat shock. B2 SINE is a crucial part of this response with abundance increasing as much as 40-fold from baseline levels ^24^. The accumulation of B2 SINE transcripts selectively downregulates RNA Pol II-dependent transcription by disrupting Pol II association with the promoter ^25,26^. When B2 SINE induction is inhibited, the heat shock response becomes dysregulated and the expression of these HKGs remains “on” ^23^. In order to circumvent the potential confound of B2 SINE regulation of RNA Pol-II HKGs, we chose to use two different RNA Pol III-dependent controls, 7SK and 5S, in parallel for normalizing both B2 SINE and ß-actin expression levels in the adult rat hippocampus. We have previously shown that B2 SINE is also a psychologically stress-sensitive transposable element that transiently generates a non-coding RNA under psychologically stressful conditions, such as acute restraint stress, or developmental stress such as maternal immune activation (MIA) ^27,28^. Under basal conditions, these elements are epigenetically silenced in rat hippocampus, but under acute restraint stress, this particular element is released from epigenetic silencing and allowed to transcribe into RNA ^29,30^. B2 RNA then colocalizes with glucocorticoid receptor to alter its transcriptional regulation, something that has been shown *in vivo* and *in vitro* ^29,31^. Others have shown that B2 RNAs have increased processing in the hippocampi of rodents with amyloid pathology ^32^. The study that follows is the first to knock down B2 SINE RNA through an siRNA GapmeR pool in primary hippocampal neurons to assess the sex differences in dendritic arborization in combination with androgen treatment.

## Results

B2 SINE RNA is an allosteric regulator of pol II-dependent HKG expression. Therefore, we measured the expression of two pol III-dependent HKGs, 5S and 7SK, in tandem with both B2 SINE and ß-actin in the adult rat hippocampus. A significant difference was observed for both B2 SINE ( t(30)=2.2358, p<.05) and ß-actin ( t(30)=11.504, p<1×10-12) normalized to 5s. B2 SINE expression was higher in males compared to females (Fig. 1A) whereas ß-actin expression was lower in males compared to females (Fig. 1B). A weak, not statistically significant negative correlation ( r(30)=.86803, p=.1961) was observed between B2 SINE and ß-actin deltaCT values (Fig. 1C)

**Figure 1.**
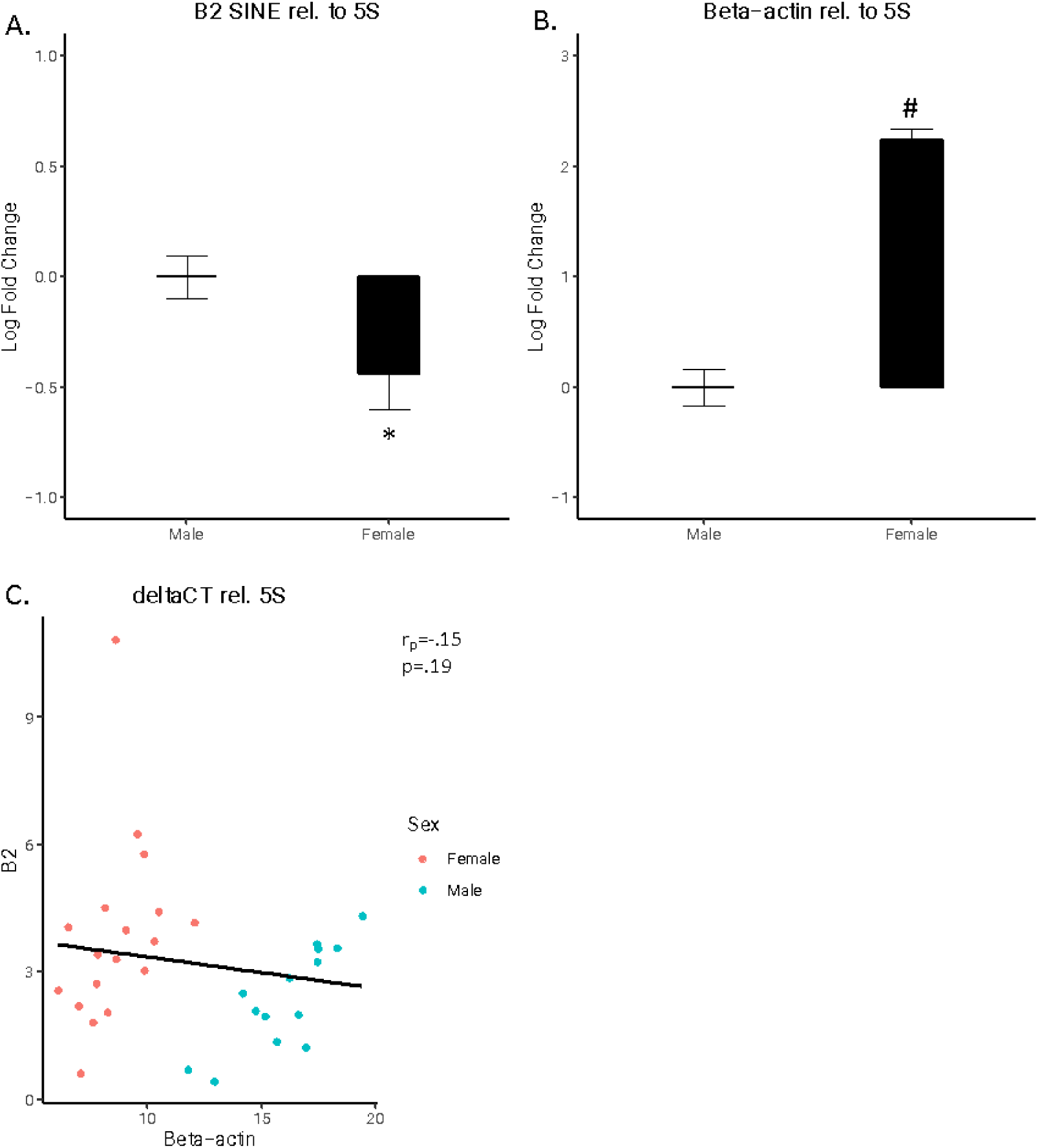
B2 SINE and Beta-actin expression relative to 5S. A. B2 SINE expression and B. Beta-actin expression. Log fold changes are shown for B2 SINE and beta-actin expression relative to 5S. **C. B2 SINE vs. beta-actin expression**. DeltaCT values for B2 SINE and beta-actin are shown relative to 5S. Males are specified by blue squares and females by red circles. The 95% confidence interval for predictions from a linear model is noted in grey. (*p<.05, #p<10-10)

A significant difference was also observed for both B2 SINE and ß-actin normalized to 7SK. Once again, B2 SINE expression was higher in males compared to females (Fig. 2A; t(30)=3.1148, p<.01) whereas ß-actin expression was lower in males compared to females (Fig. 2B; t(30)=12.37, p<1×10-13). A significant negative correlation was observed between B2 SINE and ß-actin deltaCT values (Fig. 2C; r(30)=1.91, p<.05).

**Figure 2.**
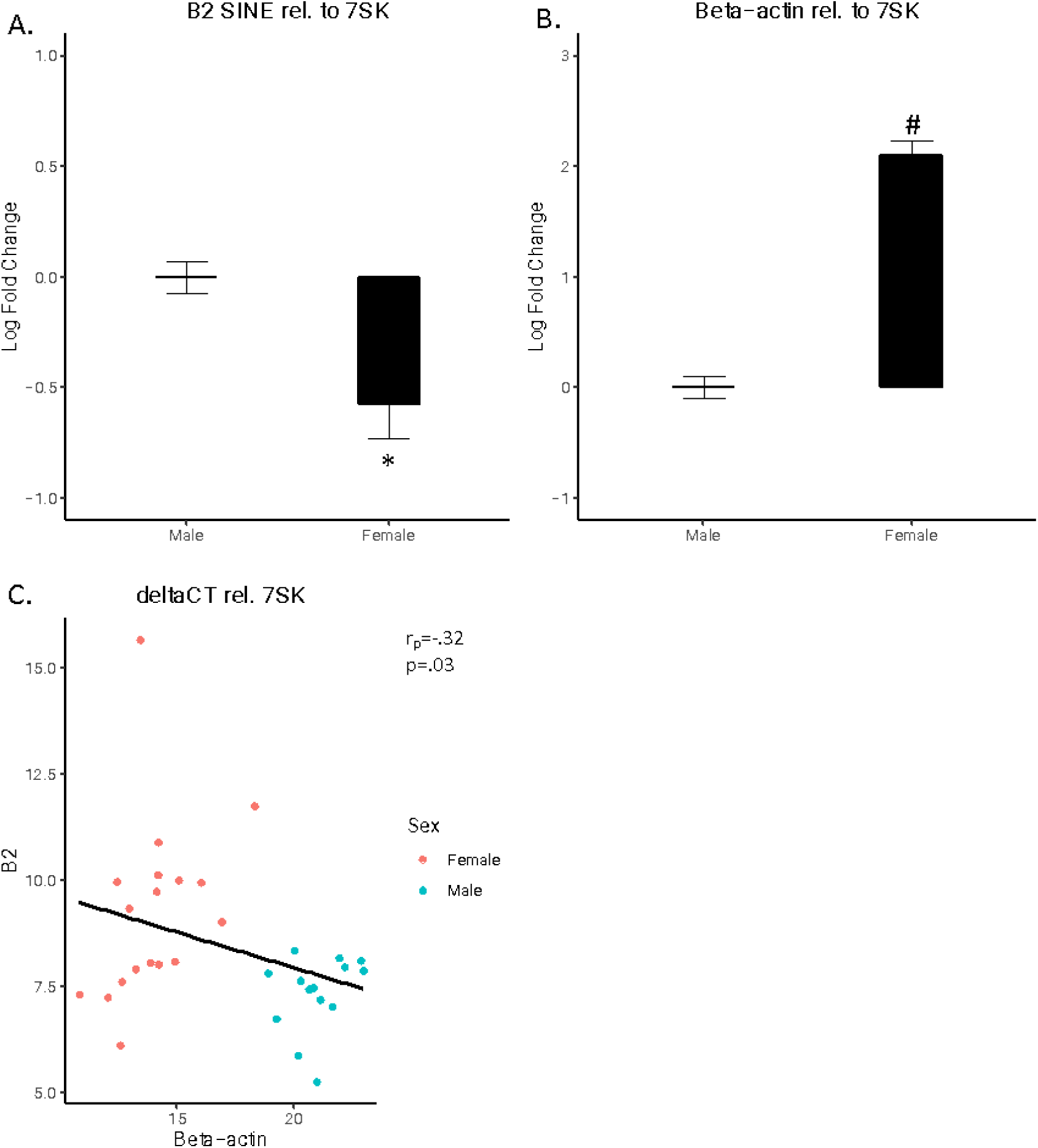
B2 SINE and Beta-actin expression relative to 7SK. A. B2 SINE expression and B. Beta-actin expression. Log fold changes are shown for B2 SINE and beta-actin expression relative to 7SK. **C. B2 SINE vs. beta-actin expression.** DeltaCT values for B2 SINE and beta-actin are shown relative to 7SK. Males are specified by blue and females by red. The 95% confidence interval for predictions from a linear model is noted in grey. (*p<.05, #p<10-10)

We tested whether a subset of B2 SINE and ß-actin expression could be used to generate a model to successfully predict sex. A principal component analysis classified the data into two distinct groups regardless of HKG used for normalization (Fig. 3A-B). Using linear discriminant analysis, we created a model using a 0.60 subset of the data for training. The models generated predicted the categorical variable, sex, without error regardless of HKG used for normalization (Fig. 3C-D).

**Figure 3.**
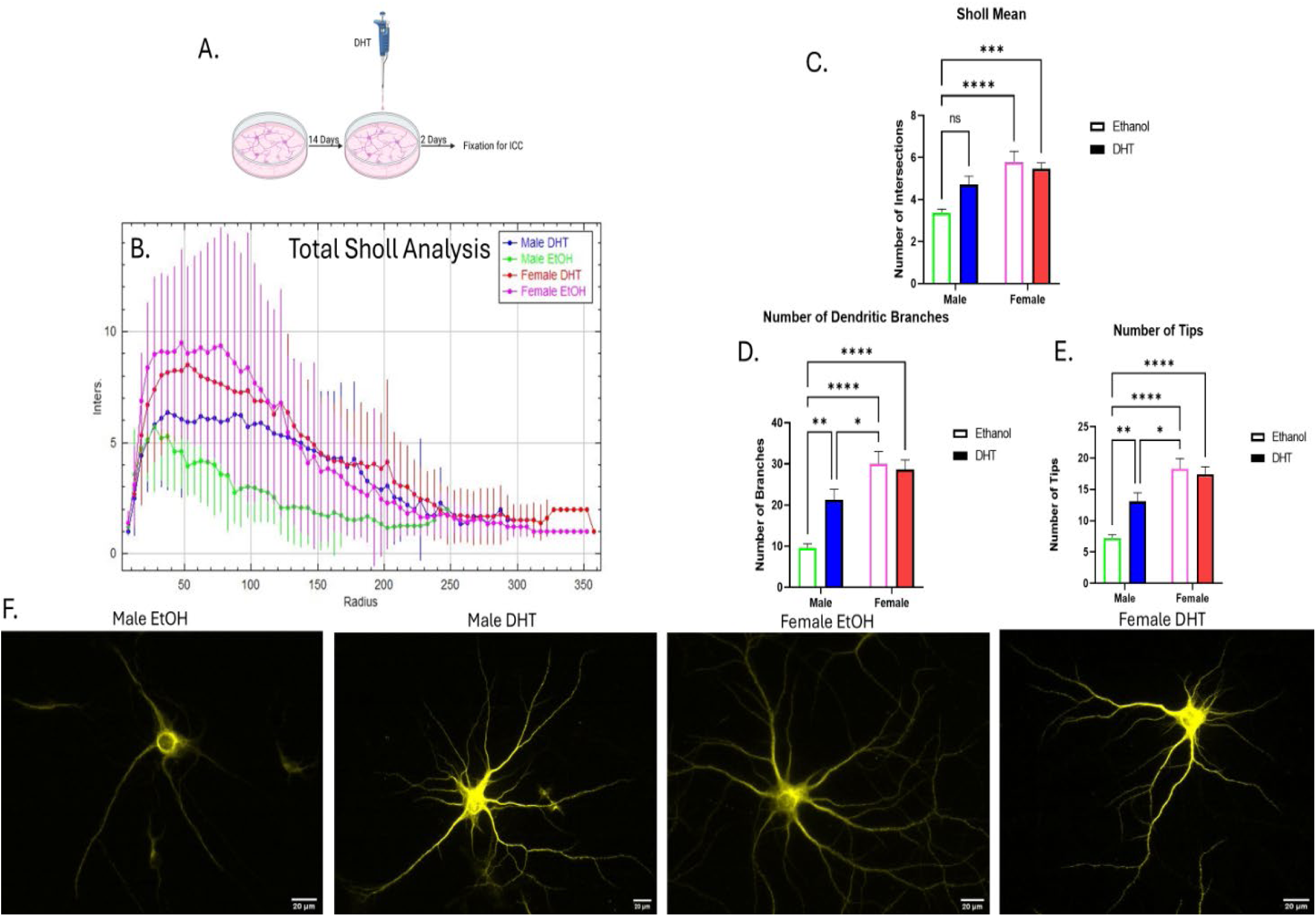
Sholl analysis of treated male and female primary hippocampal neurons. (A) Experimental paradigm for primary hippocampal culture knockdown. (B) Total Sholl Analysis for all eight groups. Data is represented as the number of intersections (Inters.) for each concentric circles radius (Radius) ± SD (Male-DHT-KD n = 50, Male-DHT-Scramble n = 45, Male-EtOH-KD n = 39, Male-EtOH-Scramble n = 42, Female-DHT-KD n = 51, Female-DHT-Scramble n = 41, Female-EtOH-KD n = 37, Female-EtOH-Scramble n = 42). (C) Sholl Mean, (D) Number of Dendritic Branches, (E) Number of Tips is shown. Data is represented as mean ± SEM (*p<0.05, **p<0.01, ***p<0.001, ****p<0.0001). (F) Representative images of male MAP2-stained neurons.

**Figure 4.**
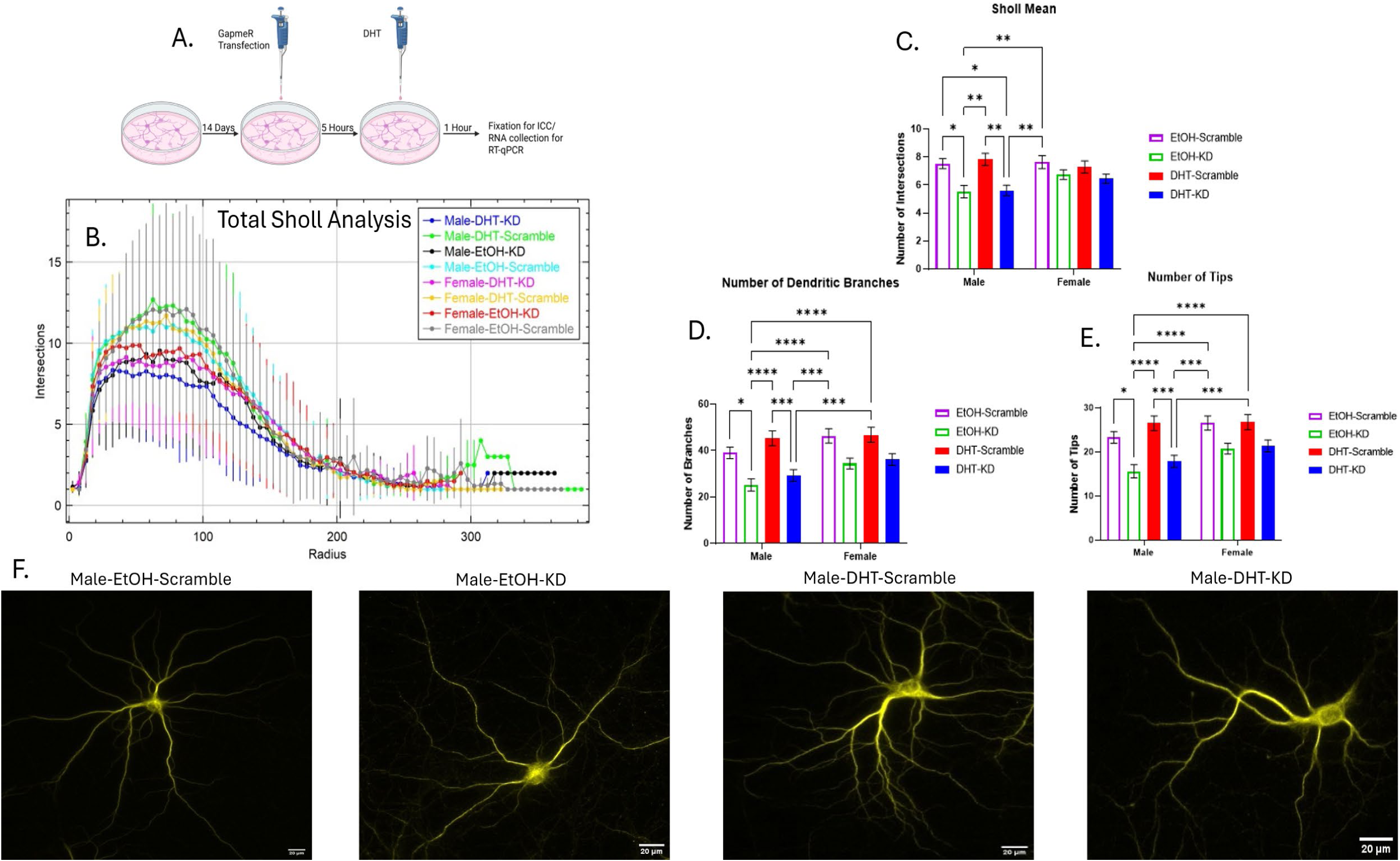
Sholl analysis of treated and knocked down male and female primary hippocampal neurons. (A) Experimental paradigm for primary hippocampal culture. (B) Total Sholl Analysis for all four groups. Data is represented as the number of intersections (Inters.) for each concentric circles radius (Radius) ± SD (Male DHT n = 27, Male EtOH n = 26, Female DHT n = 27, Female EtOH n = 26). (C) Sholl Mean, (D) Number of Dendritic Branches, (E) Number of Tips is shown. Data is represented as mean ± SEM (ns: not significant, *p<0.05, **p<0.01, ***p<0.001, ****p<0.0001). (F) Representative images of MAP2 stained neurons.

### DHT promotes dendritic complexity in male primary hippocampal neurons but not in females

To investigate the role DHT plays in dendritic arborization over the long term, DHT was added to neurons for two days, and dendritic arborization was assessed. Neurons were stained with Microtubule Associated Protein 2 (MAP2), a marker that shows up in dendrites and somas but not axons.

For Sholl mean, after a two-way Anova, there was a significant main effect of sex (η^2^ = 0.1449, p<0.0001) and an interaction effect (η^2^ = 0.04037, p<0.05). After multiple comparisons, there was a difference between the male neurons that received ethanol and both groups of female neurons (Figure 6B and 6C). However, there was no difference between male neurons treated with ethanol and males treated with DHT.

For the mean number of dendritic branches, after a two-way Anova, there was a significant main effect of Sex (η^2^ = 0.2357, p<0.0001) and Treatment (η^2^ = 0.03259, p<0.05) and an interaction effect (η^2^ = 0.05214, p<0.01). After multiple comparisons, there was a difference between male neurons that received ethanol and male neurons that received DHT, female neurons that received ethanol, and female neurons that received DHT. Finally, there was a difference between male neurons that received DHT and female neurons that received ethanol (Figure 6D).

For the mean number of tips, after a two-way Anova, there was a significant main effect of Sex (η^2^ = 0.2509, p<0.0001) and an interaction effect (η^2^ = 0.04835, p<0.01). After multiple comparisons, there was a difference between male neurons treated with ethanol and male neurons treated with DHT. Additionally, there was a difference between male neurons treated with ethanol and both groups of female neurons. Finally, there was a difference between male neurons treated with DHT and female neurons treated with ethanol (Figure 6E). Representative images of 1 neuron from each group are pictured (Figure 6F).

### Knocking down B2 SINE RNA reduces dendritic complexity in male primary hippocampal neurons but not in females

We utilized pooled B2 GapmeRs to knock down (KD) the expression of B2 SINE RNA in primary hippocampal neurons. We then treated these neurons with DHT or ethanol and measured the dendritic complexity through Sholl analysis.

For Sholl mean, after a three-way Anova, there was a significant main effect of knockdown (η^2^ = 0.07356, p<0.0001) and an interaction effect between sex and knockdown (η^2^ = 0.01290, p<0.05). After multiple comparisons there was a difference between Male-DHT-KD and Male-DHT-Scramble, Male-DHT-KD and Male-EtOH-Scramble, Male-DHT-KD and Female-EtOH-Scramble, Male-DHT-Scramble and Male-EtOH-KD, Male-EtOH-KD and Male-EtOH-Scramble, and Male-EtOH-KD and Female-EtOH-Scramble (Figure 8B and 8C).

For the number of branches, after a three-way Anova, there was a significant main effect of knockdown (η^2^ = 0.1104, p<0.0001) and a simple main effect of sex (η^2^ = 0.02507, p<0.01). After multiple comparisons, there was a difference between Male-DHT-KD and Male-DHT-Scramble, Male-DHT-KD and Female-DHT-Scramble, Male-DHT-KD and Female-EtOH-Scramble, Male-DHT-Scramble and Male-EtOH-KD, Male-EtOH-KD and Male-EtOH-Scramble, Male-EtOH-KD and Female-DHT-Scramble, and Male-EtOH-KD and Female-EtOH-Scramble (Figure 8D).

For the number of tips, after a three-way Anova, there was a significant main effect of knockdown (η^2^ = 0.1079, p<0.0001) and a simple main effect of sex (η^2^ = 0.02098, p<0.01). After multiple comparisons, there was a difference between Male-DHT-KD and Male-DHT-Scramble, Male-DHT-KD and Female-DHT-Scramble, Male-DHT-KD and Female-EtOH-Scramble, Male-DHT-Scramble and Male-EtOH-KD, Male-EtOH-KD and Male-EtOH-Scramble, Male-EtOH-KD and Female-DHT-Scramble, and Male-EtOH-KD and Female-EtOH-Scramble (Figure 8E). Representative images of 1 neuron from the male groups are pictured (Figure 8F).

Altogether, this data suggests that B2 SINE RNA is playing a positive role in dendritic complexity in males’ primary hippocampal neurons but not in females.

## Discussion

We have found that B2 SINE RNA is a novel regulator of dendritic complexity of male neurons but not female neurons in primary hippocampal culture. We have also found that DHT contributes to dendritic complexity in developing male neurons but not females. We believe this is the first instance of a transposable element’s RNA being implicated in the development of the cytoarchitecture of neurons in a sex dependent manner and the first instance illustrating that dendritic arborization is altered 2 days after a DHT surge in a sex specific manner.

B2 SINEs generate ncRNA by RNA polymerase III and have a high copy number in the mouse genome ^33,34^. Typically, these elements are repressed in cells, but in times of stress can be released from silencing and upregulated ^29–31,33–37^. There are remarkably few studies examining B2 SINE RNA in the nervous system, specifically within individual neurons. The studies that do exist focus on a stress effect ^38,39^. Very recently, it was determined that B2 SINEs act as intrinsic axon growth regulators ^40^. This study showed that B2 SINE ncRNA regulates neuronal growth specifically in nerve injury regeneration. These studies suggest that B2 SINE’s ncRNA might have diverse biological roles in neurons separate from the stress effects observed. Our study suggests that B2 SINE RNA promotes dendritic complexity in the developing hippocampus but only in male neurons.

Sex differences in the rodent brain are believed to be due in large part to the actions of gonadal hormones and their respective receptors. What is unclear, however, is the mechanism by which males’ and females’ hippocampi are differentially developed. It is believed that sexual differentiation of the rodent hippocampus starts early in life, with the organizational effects of circulating androgens and estrogens beginning *in utero* (see (Premachandran et al., 2020), for a review). We see a sex effect here of dendritic arborization in part because of androgens and not estradiol. DHT is a non-aromatizable androgen, and this hormone acts through AR. Previously, it was shown that DHT treatment contributes to a sex difference in hippocampal morphology ^14,15^, suggesting that sex differentiation of the rodent hippocampus begins early in life with the organizational effects of DHT and AR. Furthermore, unpublished data from our lab shows that B2 SINE is differentially expressed in the adult rat hippocampus, showing a significantly higher amount in males than females. Taken together with this work, this suggests male hippocampal neurons are using the higher amounts of B2 SINE ncRNA for the proper development of dendritic complexity, while females do not need this ncRNA as much.

Translationally, *ALU* SINE RNA is the closest functionally related ncRNA to B2 SINEs ^42^. B2 and *ALU* SINEs have evolved convergently to the same function in rodent and primate genomes, respectively ^43,44^. Specifically in that of Alzheimer’s pathology, where increased processing of B2 and *ALU* SINEs showed transcriptome deregulation and gene expression changes in amyloid beta neuropathology in rodents and human Alzheimer’s brains, respectively ^32,45^. Altogether, our study combined with others in the literature shows that B2 SINE ncRNA is contributing to the development of the cytoarchitecture of neurons in the brain. Our study expands on this by showing a sex effect on the development of neurons in the hippocampus.

We identified a novel sex difference in brain retrotransposon RNA expression. To our knowledge, only one previous study has identified a sex difference in retrotransposon RNA expression in the hippocampus ^46^. Further, we have shown that the hippocampal expression of B2 RNA and ß-actin, can be used to predict sex, indicating that this difference is robust. In light of the role of B2 SINE RNA during the cellular heat shock response, perhaps this retrotransposon is regulating the expression of ß-actin, which is known to vary between sexes in a variety of tissues, including the brain ^25,47,48^. In this same vein, Alu RNA has shown to similarly be critical to the heat shock response in primates acting reflectively ^49^. Alu RNA abundance has also been linked to retinal degradation in cellular models of macular degeneration and its expression is correlated with Tau pathology in the fly brain illustrating the potential for dysregulation of TEs to lead to pathologies ^50–53^. Future studies looking in these contexts may reveal sex differences in Alu RNA expression potentially conferring resistance or susceptibility to pathological phenotype. Yet the mechanisms by which B2 SINE RNA may be differentially regulated in the brain by the sexes as well as the potential implications are of profound interest for understanding TE-related pathologies. Our previous observation that B2 SINE RNA is regulated in the brain by maternal immune activation, also in a sex specific manner is salient in this context. MIA, like male sex is a significant risk factor for ASD suggests these ncRNAs may be an important target for future research into the etiology of ASD and other neurodevelopmental disorders ^27,28^. These data provide a scaffold for future studies for determining such potential effects in the rodent hippocampus and other brain regions.

## Materials and Methods

### Ethics statement

Animals were maintained in accordance with the Randolph Macon College Institutional Animal Care and Use committee. All animals were fed standard chow *ad libitum* and kept on a 12:12 light cycle in standard polycarbonate laboratory cages (48cm x 26cm x 21cm).

### Animals

Thirty-two male and female Long Evans rats were reared in-house from eight different mothers. Rats were weaned at post-natal day (PND) 21. On PND 60 18 females and 14 males, were anesthetized with 1ml halothane and upon determination that the animal was nonresponsive, it was sacrificed via rapid decapitation. Brains were dissected and hippocampi extracted prior to being flash frozen. For primary hippocampal culture, post-natal Sprague Dawley rats at day 0 of age were used.

### Primary Hippocampal Culture

Post-natal Sprague Dawley rats at day 0 of age were used to generate primary hippocampal culture as previously described with minor modifications ^54,55^. Rats were sexed by measuring the anogenital distance. Hippocampi were aseptically dissected bilaterally from pups and placed into HBSS (Hanks Balanced Salt Serum, HEPES, and Antibiotic-Antimycotic). Hippocampi from males and females were placed in their corresponding tubes based on sex. Combined hippocampi were then dissociated with trypsin and DNase I and then triturated. Cell viability was determined by the trypan blue exclusion method. Dissociated cells were then added at a concentration of 4 x 10^5^ to etched poly-L coverslips in 6-well plates for 2 to 4 hours in plating medium (Minimum essential medium, charcoal stripped fetal bovine serum, glucose, and pyruvic acid) in a humidified 37°C incubator with 5% CO_2_. Once cells were viable and attached, plating media was aspirated and Neurobasal+ (Neurobasal media, L-Glutamine, Antibiotic-Antimycotic, B-27 supplement) was added to wells containing coverslips. Every 2-3 days, 50% of the media was replaced with fresh media until days *in vitro* (DIV) 14. For the DHT experiment, on DIV 14, 100 nM dihydrotestosterone (DHT) or vehicle (ethanol) was added to cultures for 48 hours. On DIV 16 immunocytochemistry staining was started for analysis of dendritic arborization. For the knockdown experiment, GapmeRs were added on DIV 14 for 5 hours, and DHT was added for 1 hour, after which immunocytochemistry staining was started.

### Immunocytochemistry

Immunocytochemistry was performed either 1 hour after DHT or 48 hours after treatment. Cells on coverslips were fixed with 4% formaldehyde and subsequently blocked in phosphate-buffered saline (PBS) supplemented with 5% normal goat serum and 0.3% Triton X-100. Hippocampal neurons were stained with a primary polyclonal antibody against Map 2 (4542S) (Cell Signaling) (1:1600). Primary antibody was detected with goat antirabbit-IgG AlexaFluor 555 (Cell Signaling) (1:1000). Nuclei were counterstained and coverslips mounted with ProLong gold antifade mounting medium with DAPI (Cell Signaling). Images were captured on a Leica DM5500 B confocal microscope (Leica) equipped with a K3M camera (Leica) with a 20x objective. All images were taken randomly near the center of the coverslip. Between N=25 and 50 neurons were analyzed for each group. Images were captured and transferred for neuronal tracing.

### Neuron Tracing

Tracing of primary hippocampal neurons was semi-automated. 12-bit images of primary hippocampal culture neurons were traced using Simple Neurite Tracer (SNT) ^56^ which is a part of the Neuroanatomy plugin for Fiji ^57^. Tracing was performed blind to the group membership of individual neurons.

### Branch Number and Sholl Analysis

Sholl, branch number and branch tip analysis was performed using SNT using default settings. A linear plot containing the calculated best-fit polynomial and a detailed table were generated and saved then merged and compared using a script from SNT. The mean, standard deviation, and number were then brought over to Prism for statistical analysis.

### LNA GapmeRs

To knock down the expression of B2 SINE RNA, we used a pool of Locked Nucleic Acids (LNA) GapmeRs adapted from ^32,58^ with minor modifications. Anti-sense Oligonucleotides (ASO) were utilized to bind to B2 SINE RNAs to elicit RNase-H mediated degradation. The ASOs were synthesized into a pool of 4 unique sequences. Each sequence was synthesized as single-stranded DNA bases, each flanked with three 2-O-methyl modified RNA bases linked by a phosphorothioate backbone (Integrated DNA Technologies). The sequence of scrambled B2 GapmeR is synthesized as single-stranded 2-O-methyl modified RNA bases linked by a phosphorothioate backbone. The sequences of the GapmeRs are listed in supplemental methods.

### Transfection of LNA GapmeRs

LNAs were provided as dried down flakes and subsequently reconstituted with ultrapure nuclease-free water (ThermoFisher). On the day of LNA transfections, DIV 14, a pool was made by mixing 10μM of each LNA GapmeR into one tube. The pool was then combined with RNAiMAX reagent (Lipofectamine, ThermoFisher), and neurobasal+ media and incubated for 5 minutes at room temperature. LNAs were then added dropwise to the cells to a final amount of 25 pmol per cover slip. Control cells received a scrambled version instead. 10 nM DHT was added 5 hours after transfection for 1 hour. Successful transfection was verified by detection of B2 SINE RNA with RT-qPCR.

### RNA extraction and cDNA preparation

RNA was extracted using RNeasy Lipid Tissue Mini kit according to the manufacturer’s protocol (Qiagen). RNA was then analyzed via Nanodrop2000 for purity and concentration. Residual gDNA was removed and cDNA then prepared using random hexamers with the QuantiTect Reverse Transcription kit according to the manufacturer’s protocol (Qiagen).

### RT-qPCR

Sybr green master mix or TaqMan master mix was used to determine relative expression by the deltaCt method. Primers used targeting the consensus sequences for *rattus norvegicus* B2 SINE and ß-actin were as previously described ^29^. *Rattus norvegicus* 7SK snRNA and 5S rRNA primers were built against the refSeq RNA sequence using IDT’s primer design tool and verified for specificity using Primer BLAST. Sequences are listed in supplemental methods.

### Statistical Analysis

All statistical analysis done using R or GraphPad Prism. A student’s t-test was used to assess differences in B2 and ß-actin expression between the sexes. Pearson’s correlation test was used to determine correlations between B2 and ß-actin expression. The “factoextra” and “cluster” R packages were used for PCA analysis. The “MASS” and “klaR” R packages were used for the linear discrimination analysis (sampling with replacement sixty percent of the data set for training)^59,60^. Statistical significance for dendritic analysis after DHT was determined by a two-way ANOVA with a Tukey’s correction for multiple comparisons. Statistical significance for dendritic analysis after RNA knockdown was determined by three-way ANOVA with a Tukey’s correction for multiple comparisons. All data are represented as the mean ± standard error of the mean, except for the Sholl mean line chart, where data is represented as the mean ± standard deviation. Statistical significance threshold was set at p < 0.05.

## Data availability

The data analyzed during the current study are available from the corresponding author on reasonable request.

Supplementary Information is available for this paper.

Correspondence and requests for materials should be addressed to Troy A. Richter, PhD. Reprints and permissions information is available at www.nature.com/reprints.

## Acknowledgements

We wish to thank Drs. Kelly Lambert and Molly Kent for providing brain tissue for this study.

## Author contributions

TR, EO, EP, AB, HL, acquired data. TR, EO, AB, GG and HL analyzed and interpreted the data. TR, EP AB, HL, SZ and RH designed the experiments. TR and AB wrote the manuscript and all authors contributed to manuscript revision.

