## Supplemental Information for "Sex Differences in B2 SINE RNA Expression and its Role in Hippocampal Development"

**Supplementary Data:**

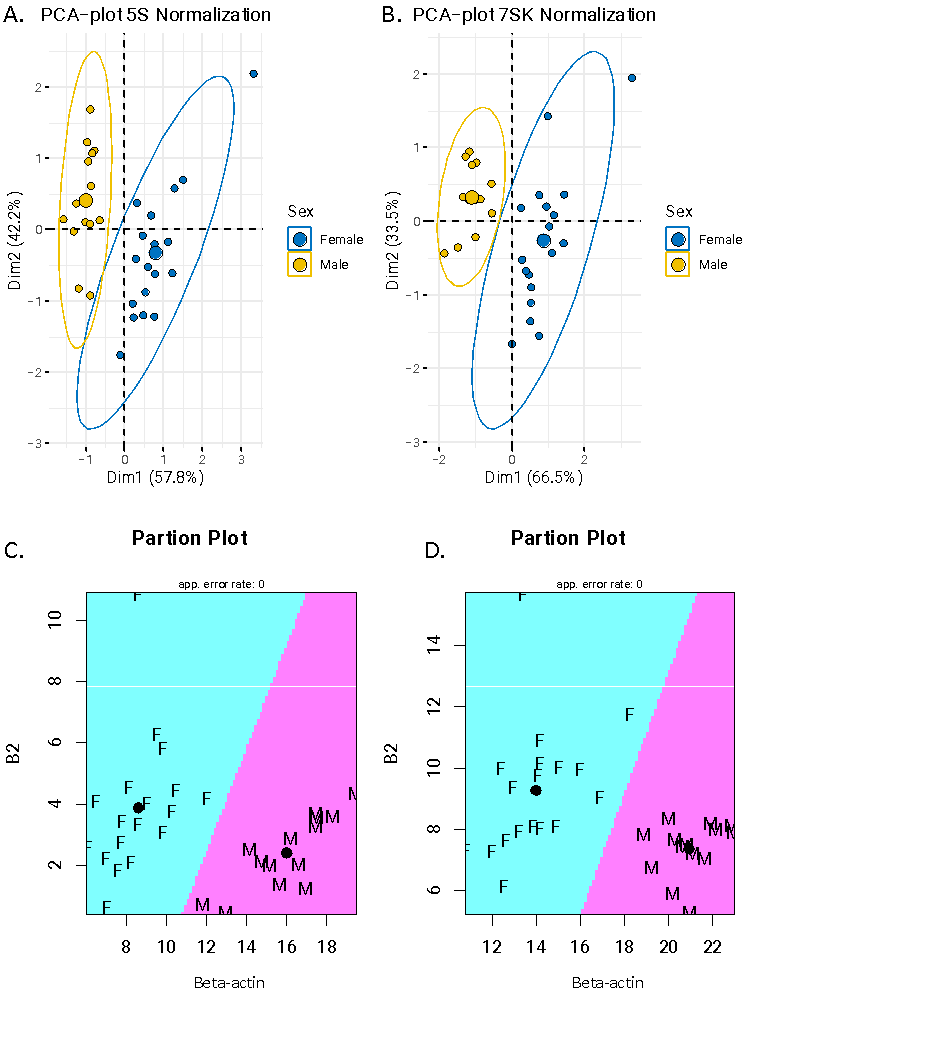

**Figure S1. B2 SINE and beta-actin expression predictive models.** A. Principal component analysis plot using expression values normalized to 5S and B. to 7SK PCA successfully segregates data into two clusters. C. Partition plot from linear discriminant analysis normalized to 5S and D. to 7SK. The models created from a subset of the data correctly predicts sex.

**Figure S2. LNA GapmeR pool targets and knocks down B2 SINE RNA in primary hippocampal cells.** Expression of B2 SINE RNA after either GapmeR pool or scrambled vector transfection. Data is represented as mean fold change over 7sk RNA ± SEM (n = 3/group) (**p<0.01).
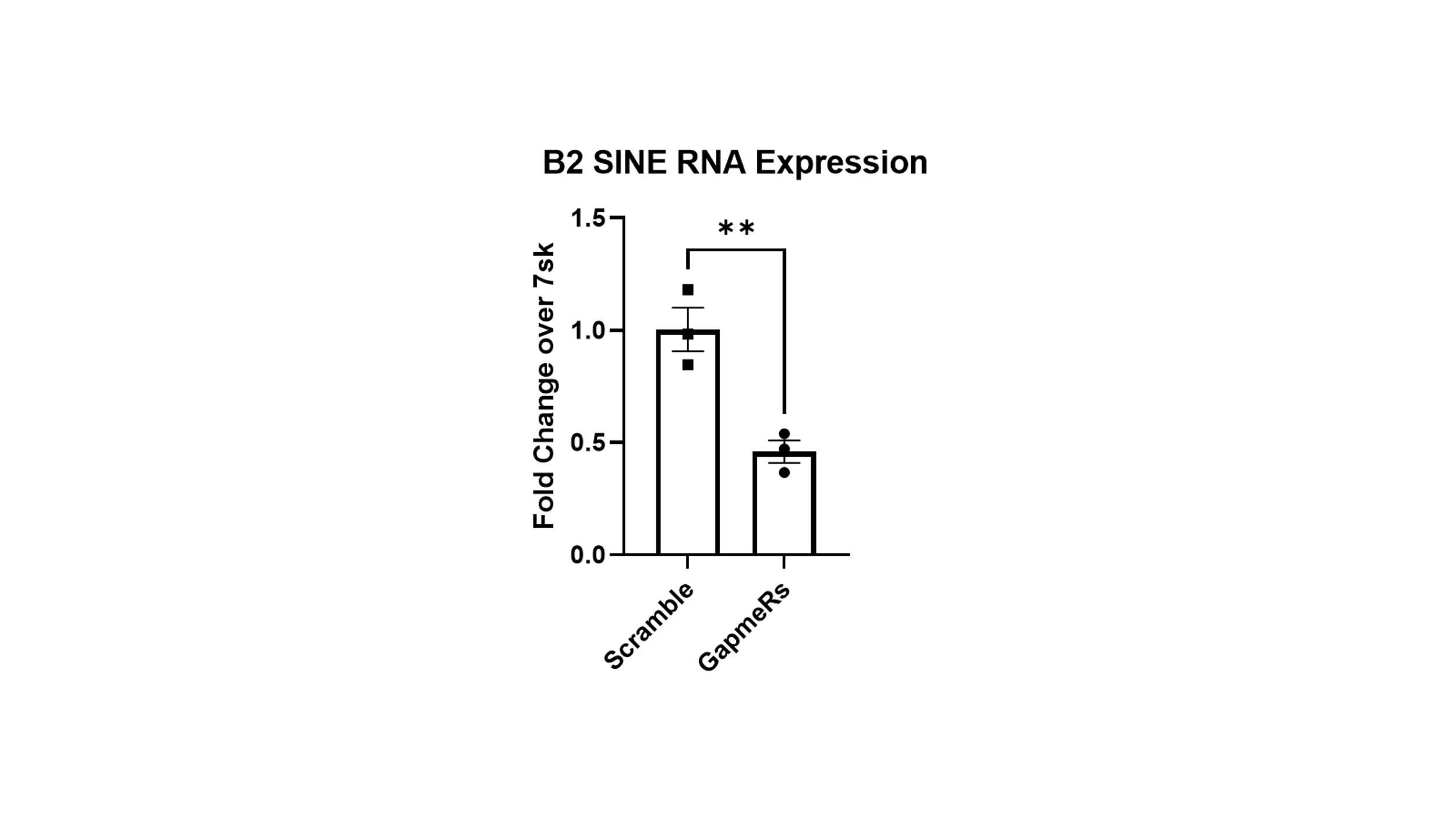

In order to assess if the pooled B2 GapmeRs successfully knocked down B2 SINE RNA in primary hippocampal neurons after 6 hours, we transfected a subset of neurons for 6 hours, collected RNA, and tested for presence by RT-qPCR. We found a significant difference between the neurons transfected with the GapmeR pool and neurons transfected with a scrambled version (Figure 7). This shows that B2 SINE RNA is depleted after transfection with GapmeRs for 6 hours.

**Supplementary Methods:**

Primer sequences and GapmeR sequences are as follows. The sequences of B2 GapmeRs are 5′-UUC(dA)(dA)(dA)(dT)(dC)(dC)(dC)(dA)(dG)(dC)(dA)(dA)(dC)(dC)(dA)(dC)(dA)(dT)(dG)(dG)(dT)(dG)(dG)(dC)(dT)(dC)(dA)(dC)(dA)ACC-3′; 5′-AGU(dT)(dC)(dA)(dA)(dA)(dT)(dC)(dC)(dC)(dA)(dG)(dC)(dA)(dA)(dC)(dC)(dA)(dC)(dA)(dT)(dG)(dG)(dT)(dG)GCU-3′; 5′-GAG(dT)(dT)(dC)(dA)(dA)(dA)(dT)(dC)(dC)(dC)(dA)(dG)(dC)(dA)(dA)(dC)(dC)(dA)(dC)AUG-3; 5′-AGC(dA)(dA)(dC)(dC)(dA)(dC)(dA)(dT)(dG)(dG)(dT)(dG)(dG)(dC)(dT)(dC)(dA)(dC)(dA)ACC-3′. The sequence of scrambled B2 GapmeR is 5’-CGGUGUGUGUAUCAUUCUCUAGUGU-3’. RN_7SK: Fwd 5’-TCGGTCAAGGGTATACGAGTAG-3’ Rev 5’- TTTGGATGTGTCTGGAGTCTTG-3’ RN_5S: Fwd 5’- CGTCTGATCTCGGAAGCTAAG-3’ Rev 5’- CCTACAGCACCCGGTATTC-3’. B2 FWD 5′AGATGGCTCAGCGGTTAAGA-3’; B2 REV 5′-GACACACCAGAAGAGGGTATCA-3’
